# Evaluation of commercial qPCR kits for detection of SARS-CoV-2 in pooled samples

**DOI:** 10.1101/2020.05.28.120667

**Authors:** Vlad Petrovan, Virgil Vrajmasu, Paula Dimon, Mihaela Zaulet

## Abstract

Due to the current pandemic, global shortage of reagents has drawn interest in developing alternatives to increase the number coronavirus tests. One such alternative is sample pooling. Here we compared commercial kits that are used in COVID-19 diagnostics, in terms of sensitivity and feasibility for use in pooling. We showed that pooling of up to 60 samples did not affect the efficiency of the kits. Also, the RNA dependent RNA polymerase (RdRp) is a more suitable target in pooled samples than the Envelope (E) protein. This approach could provide an easy method of screening large number of samples and help adjust different government regulations.

## Introduction

The recent emergence of the novel severe acute respiratory syndrome coronavirus 2 (SARS-CoV-2) in December 2019 from Wuhan, China, has caused more than 5 million cases with an estimate of more than 300 000 deaths associated with coronavirus disease (COVID-19) (*1*). Clinical manifestation of COVID-19 infection is variable ranging from asymptomatic to severe disease, with symptoms including respiratory distress, fever, cough, dyspnea, and viral pneumonia (*2*). Since there is currently no targeted therapeutics against SARS-CoV-2 and clinical manifestations are not disease specific, diagnostic screening and implementation of strict biosecurity measures issued by governments are limiting the spread of the disease (*3*). The most widely used molecular method approved by the World Health Organization (WHO) and Centers for Disease Control and Prevention (CDC) to detect SARS-CoV-2 is the real-time reverse transcription-polymerase chain reaction (qRT-PCR) (*4*). In the case of a public health emergency, most of diagnostic laboratories worldwide can rely on this technology to routinely perform diagnostic services until standardized tests are widely available. Different PCR assays were rapidly developed to target the ORF1a/b, ORF1b-nsp14, RdRp, S, E, or N gene of SARS-CoV-2 and other related betacoronaviruses, such as the closely related SARS-CoV (*3, 5*). The majority of qPCR tests are using different sample matrixes, represented by either swabs or sputum, since they contain relatively high titer virus, due to the initial viral replication in the upper respiratory tract (*6*). However, the global need for a new surveillance approach reflects the requirement to adapt to the increased demand of number of molecular tests to adjust the lockdown policies.

Diagnostic pooling has been already shown to be effective both in veterinary medicine, detecting various diseases induced by swine influenza, African swine fever virus or foot-and-mouth disease virus (*7*; *8*; *9*) and in human medicine for human immunodeficiency virus (HIV) and other transfusion-transmittable diseases (*10; 11*). Recently, the same approach showed encouraging results for SARS-CoV-2 when a pool of up to 7 samples was used before the extraction and up to 60 samples could be pooled after (*12*).

Therefore, our main goal was to evaluate and compare some commercial kits currently used for COVID-19 diagnostics, using the sample pooling approach. We also showed that the high sensitivity of RNA dependent RNA polymerase (RdRp) compared to other targets, for detection of SARS-CoV-2 in pooled samples.

## Materials and methods

### Sample collection and processing

Samples included in this study consisted of swabs collected from both nostrils and the throat or sputum, following the WHO recommendations by healthcare providers and sent to the Molecular laboratory VetWork Diagnostics, Tulcea, Romania and Personal Genetics, Bucharest, Romania. A volume of 200 ul of the transport swab buffer was mixed with 500 ul lysis buffer and RNA was extracted using Power Prep Viral DNA/RNA Extraction kit, (Kogene Biotech, Seoul, Republic of Korea). Sputum samples were mixed with equal volume of PBS and processed as described above. We obtained samples tested between April 20-27, 2020. Samples collected from 24 confirmed COVID-19 patients were extracted, aliquoted and stored at –80°C until use. Negative samples were collected from 60 healthy volunteers with no COVID-19 associated symptoms.

### RNA extraction

Extraction of RNA was performed with Power Prep Viral DNA/RNA Extraction Kit (Kogene biotech, Seoul, Korea) according to the manufacturer’s instruction. After lysis, samples were incubated at room temperature for 10 minutes, after incubation 700 μl of binding buffer was added, followed by vortex and spin. Supernatant was passed through the Binding Column. The columns were washed two times with 500 μl Wash Buffer A first and Wash Buffer B second. At the final 50 μl of Elution Buffer was added and incubated for 1 minute at room temperature, followed by a spin at 13.000 rpm for one minute and eluted RNA collected.

### Real Time PCR analysis

The RNA samples were treated with PowerCheck 2019-nCoV Real-time PCR Kit (Kogene biotech, Seoul, Korea), COVID-19 PCR Diatheva Detection Kit (Diatheva, Cartoceto, Italy) and 2019 nCoV CDC EUA KIT (IDT DNA, Coralville, Iowa, USA) mixed with FastGene Probe One Step Mix (Nippon Genetics Europe Gmbh, Duren, Germany). The high specificity of the commercial assays is based on the unique sequence of the primers specific for the SARS-CoV-2 genomic sequence along with optimal PCR conditions used for amplification. This assay does not cross-react with other respiratory viruses.

PowerCheck 2019-nCoV Real-time PCR Kit provides testing solution for Wuhan coronavirus, specifically targeting the E gene for beta Coronavirus and the RdRp gene for 2019-nCoV in bronchoalveolar lavage fluid, sputum, nasopharyngeal swab and oropharyngeal swab. Kit contains: RT-PCR mix, Primer/Probe mix 1 (E gene), Primer/Probe mix 2 (RdRp gene), control 1 (E gene) and control 2 (RdRp gene). Protocol for PowerCheck 2019-nCoV Real-time PCR Kit is: 11 ul are mixed with 4 ul of each primer/probe mix and 5 ul of template RNA with a total volume of 20 ul.

COVID-19 PCR Diatheva Detection Kit allows the qualitative detection of SARS-CoV-2 RNA in upper and lower respiratory samples, is a One-step real-time reverse transcription multiplex assay bassed on fluorescent-labelled probe used to confirm the presence of RdRp gene and E gene. The assay includes also RNase P target as internal positive control (IC). Protocol for COVID-19 PCR Diatheva Detection Kit used is: 5 ul of mix 1 are mixed with 0.625 ul of mix 2, 9.375 ul of primer/probe mix and 5 ul of RNA template with a total volume of 20 ul.

2019-nCoV CDC EUA Kit mixed with Fast Gene Probe One Step Mix allows detection of N1 gene, N2 gene and RNase P gene. Protocol for 2019-nCoV CDC EUA Kit mixed with Fast Gene Probe One Step Mix used is: 10 ul of Fast Gene Probe One Step mix, 1 ul of Fast Gene Scriptase, 1.5 ul of each primer/probe 2019 n-CoV CDC EUA Kit, 2.5 ul ultrapure water and 5 ul of RNA template with a total volume of 20 ul.

PCR reactions were performed on a Real-time PCR System (7500 Real-time PCR System, Applied Biosystems, Thermo Fisher Scientific, Foster City, CA, USA) following the programs: reverse transcription for 30 minutes at 50°C, initial denaturation 10 minutes at 95° C, 40 cycles of denaturation 15 seconds at 95° C followed by an extension of 60 seconds at 60° C for PowerCheck 2019-nCoV Real-time PCR Kit; revers transcription for 30 minutes at 48°C, initial denaturation 10 minutes at 95° C, 50 cycles of denaturation 15 seconds at 95° C followed by an extension of 30 seconds at 58° C for COVID-19 PCR Diatheva Detection Kit; revers transcription for 10 minutes at 45°C, initial denaturation 2 minutes at 95° C, 40 cycles of denaturation 5 seconds at 95° C followed by an extension of 30 seconds at 55° C for 2019-nCoV CDC EUA Kit mixed with Fast Gene Probe One Step Mix. Positive to negative cutoff was set at a Ct > 40 for all the kits assayed.

### Ethical considerations

The study was conducted as part of surveillance program for COVID-19 implemented by the Romanian government, with no disclosure regarding name, physical, economic, cultural, social status of the patients, therefore did not require individual patient consent or ethical approval.

## Results

### 1. Evaluation of commercial SARS-CoV-2 qPCR kits

To determine the analytical sensitivity of the COVID-19 commercial assays used in Romanian hospitals (PowerCheck Kogene 2019-nCoV, COVID −19 PCR Diatheva Detection Kit and 2019-nCoV CDC EUA), we first evaluated their limit of detection (LOD) using the positive controls provided by the kit. Assay reproducibility was tested in duplicate using 10-fold serial dilutions of the controls and intra- and inter-assay variability evaluated for each dilution point in triplicate on two different PCR machines. The LOD of RdRp and E (end point at 10^−7^ fold dilution) was similar between the assays from Kogene and Diatheva. However, for the CDC EUA kit, LOD for N1 and N2 was 2 log unit lower (10^−4^ fold dilution) than that of E and RdRp. Therefore, for the future experiments, we decided to use Kogene and Diatheva kits.

### 2. Evaluation of different clinical specimens collected from COVID-19 infected patients

Samples were collected from patients with laboratory-confirmed COVID-19, ranging from 22 to 80 years old and consisted of swabs (from throat and/or nostrils) or sputum. Clinical presentation was either asymptomatic or mild in 14/24 patients; the rest of 10 patients either with moderate or severe COVID-19 outcome associated with co-morbidities (Table 2). Seven patients were tested negative by PCR for the initial screening, but were positive when the PCR was repeated after 2 weeks interval. Mild or asymptomatic patients did not have any other co-morbidities, and clinical signs were limited to either fever, cough and/or shortness of breath, as shown in Table 2. PCR results revealed an average Ct value for E gene of 25.78±1.16 and for RdRp, 26.05±1.18, with no variation between sample matrix used (either nasal/throat swabs or sputum).

**Table 1.**
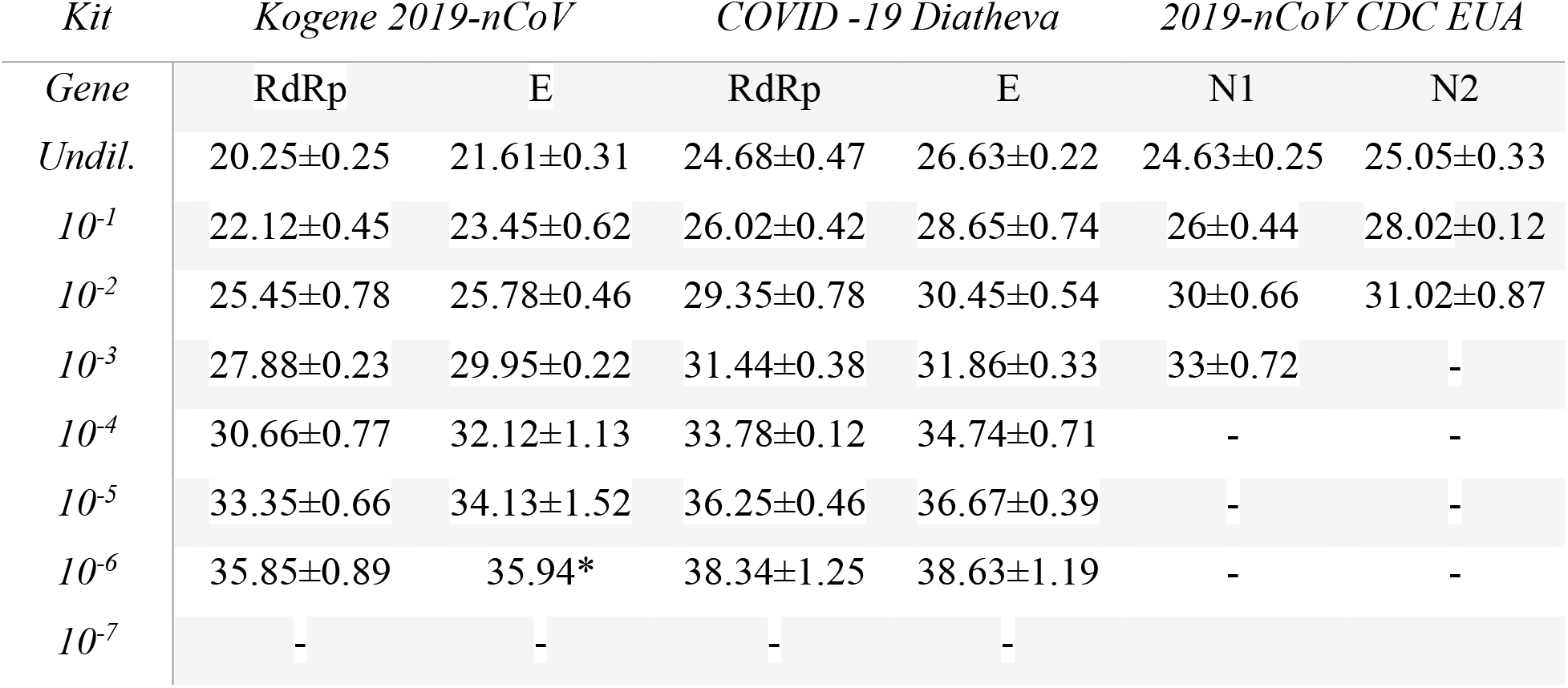
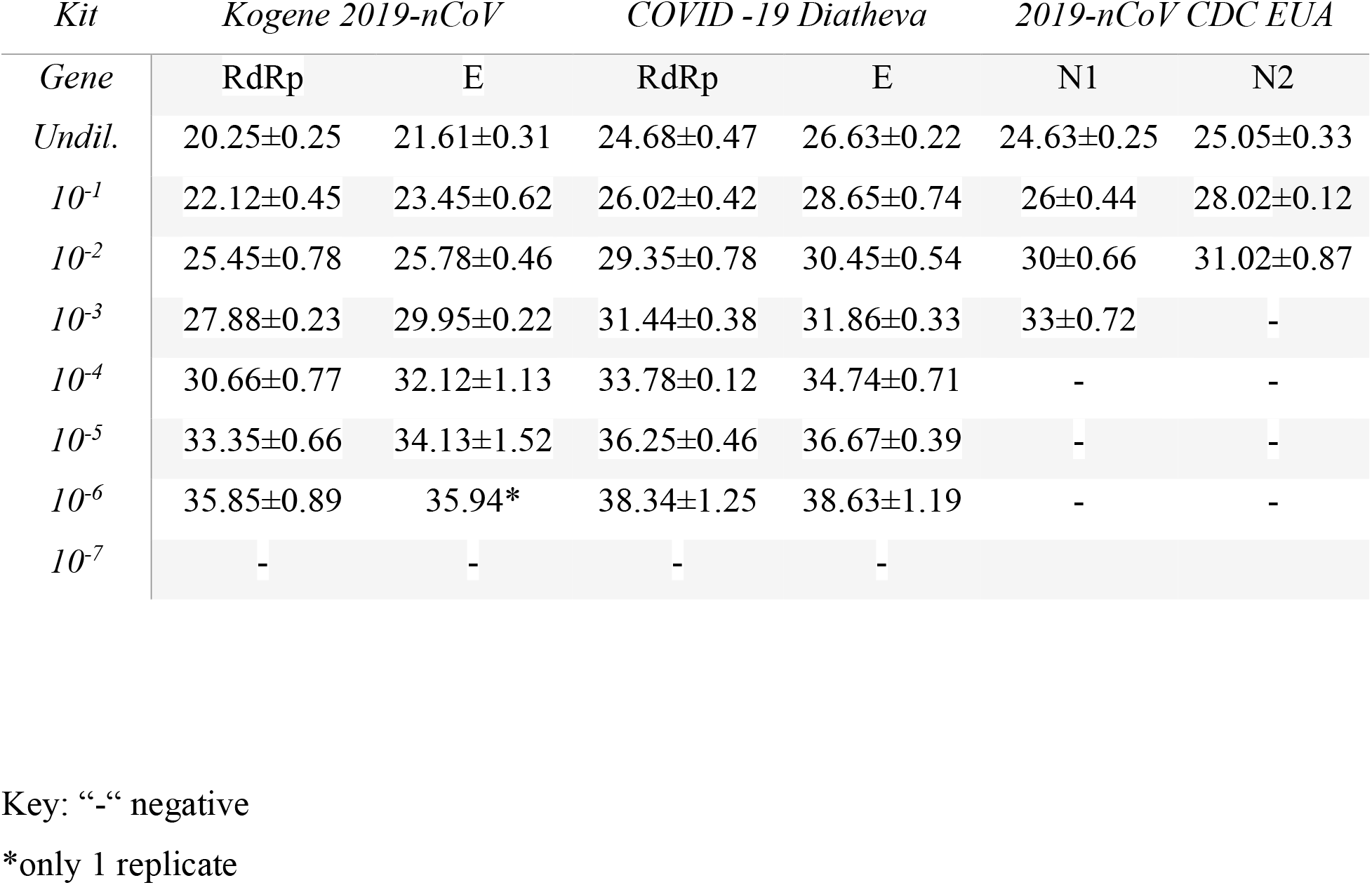
Limit of detection using the positive controls provided by the manufacturer of three commercial kits. Results are presented as average cycle threshold (Ct) of two independent experiments ± standard deviation (SD).

**Table 2.**
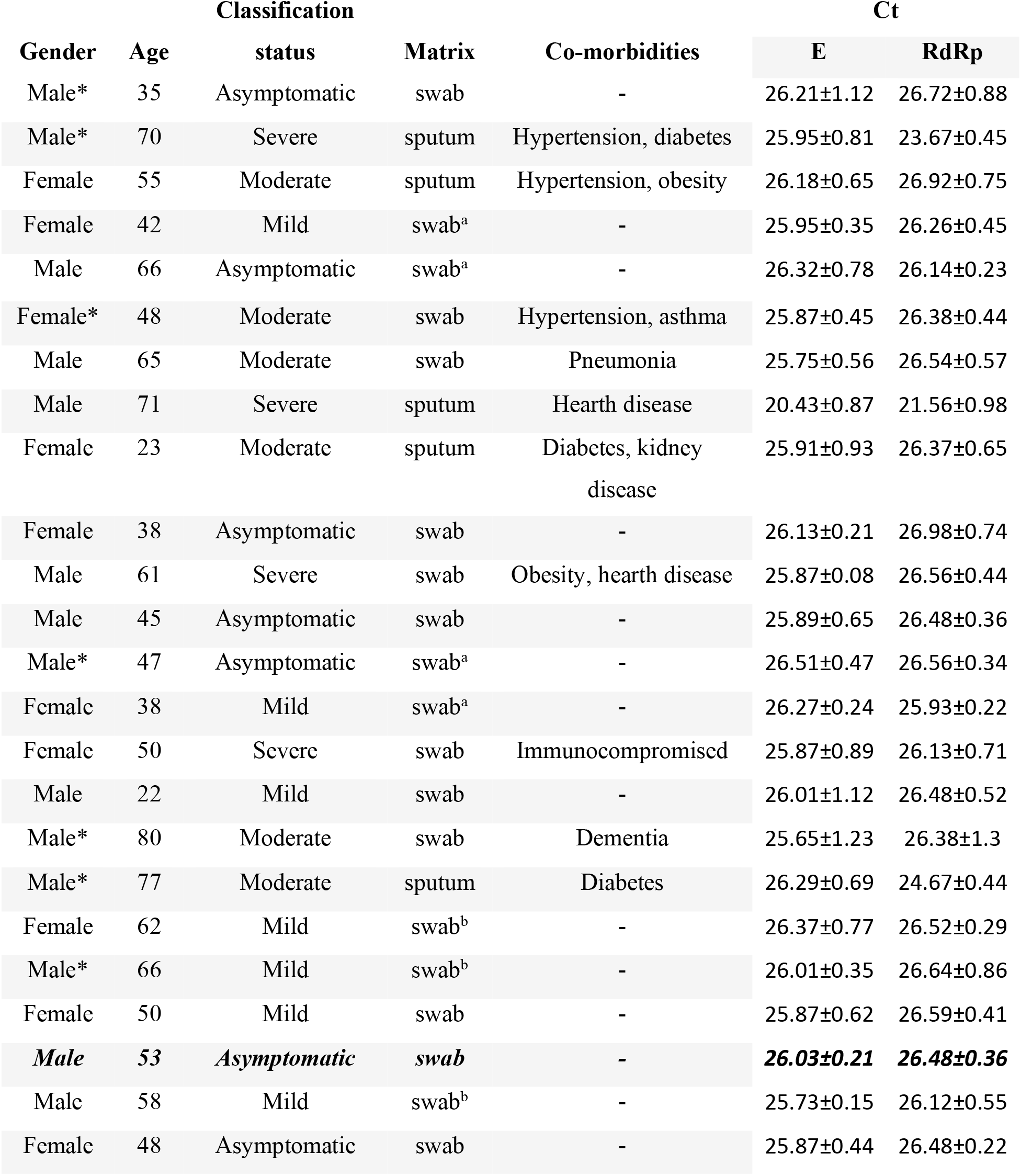
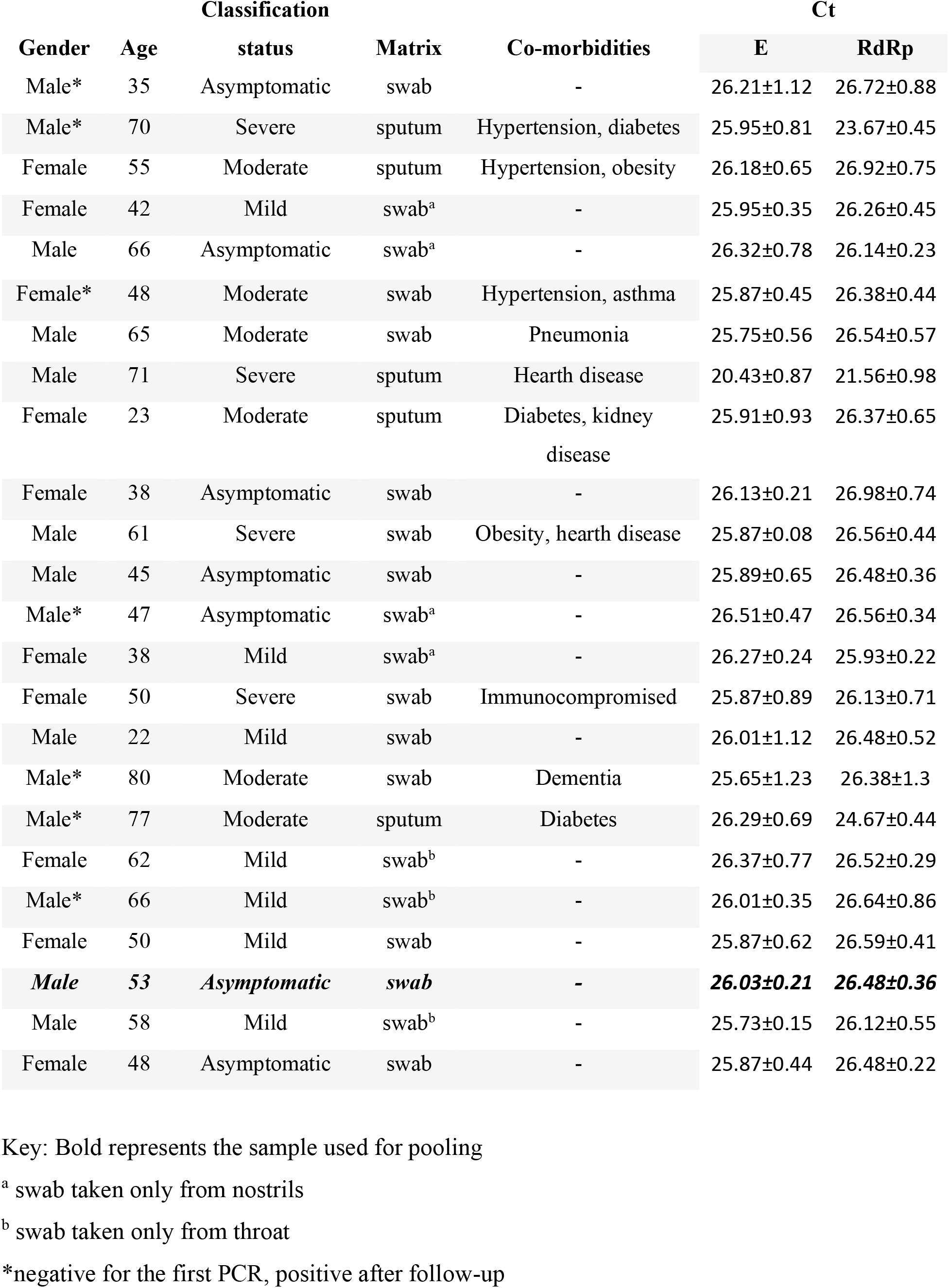
Different sample matrixes collected from COVID-19 confirmed patients.

| Gender | Age | Classification |  |  | Ct |  |
| --- | --- | --- | --- | --- | --- | --- |
|  |  | status | Matrix | Co-morbidities | E | RdRp |
| Male* | 35 | Asymptomatic | swab | - | 26.21±1.12 | 26.72±0.88 |
| Male* | 70 | Severe | sputum | Hypertension, diabetes | 25.95±0.81 | 23.67±0.45 |
| Female | 55 | Moderate | sputum | Hypertension, obesity | 26.18±0.65 | 26.92±0.75 |
| Female | 42 | Mild | swab<sup>a</sup> | - | 25.95±0.35 | 26.26±0.45 |
| Male | 66 | Asymptomatic | swab<sup>a</sup> | - | 26.32±0.78 | 26.14±0.23 |
| Female* | 48 | Moderate | swab | Hypertension, asthma | 25.87±0.45 | 26.38±0.44 |
| Male | 65 | Moderate | swab | Pneumonia | 25.75±0.56 | 26.54±0.57 |
| Male | 71 | Severe | sputum | Hearth disease | 20.43±0.87 | 21.56±0.98 |
| Female | 23 | Moderate | sputum | Diabetes, kidney disease | 25.91±0.93 | 26.37±0.65 |
| Female | 38 | Asymptomatic | swab | - | 26.13±0.21 | 26.98±0.74 |
| Male | 61 | Severe | swab | Obesity, hearth disease | 25.87±0.08 | 26.56±0.44 |
| Male | 45 | Asymptomatic | swab | - | 25.89±0.65 | 26.48±0.36 |
| Male* | 47 | Asymptomatic | swab<sup>a</sup> | - | 26.51±0.47 | 26.56±0.34 |
| Female | 38 | Mild | swab<sup>a</sup> | - | 26.27±0.24 | 25.93±0.22 |
| Female | 50 | Severe | swab | Immunocompromised | 25.87±0.89 | 26.13±0.71 |
| Male | 22 | Mild | swab | - | 26.01±1.12 | 26.48±0.52 |
| Male* | 80 | Moderate | swab | Dementia | 25.65±1.23 | 26.38±1.3 |
| Male* | 77 | Moderate | sputum | Diabetes | 26.29±0.69 | 24.67±0.44 |
| Female | 62 | Mild | swab<sup>b</sup> | - | 26.37±0.77 | 26.52±0.29 |
| Male* | 66 | Mild | swab<sup>b</sup> | - | 26.01±0.35 | 26.64±0.86 |
| Female | 50 | Mild | swab | - | 25.87±0.62 | 26.59±0.41 |
| **Male** | **53** | **Asymptomatic** | **swab** | **-** | **26.03±0.21** | **26.48±0.36** |
| Male | 58 | Mild | swab<sup>b</sup> | - | 25.73±0.15 | 26.12±0.55 |
| Female | 48 | Asymptomatic | swab | - | 25.87±0.44 | 26.48±0.22 |

Key: Bold represents the sample used for pooling
<sup>a</sup> swab taken only from nostrils
<sup>b</sup> swab taken only from throat
\*negative for the first PCR, positive after follow-up

### 3. Sample pooling and comparative performance of targets

In order to test the pooling approach, we decided to use a sample with an average Ct value of 26.25±2.1 to spike into seven negative pools containing equal volumes (200 ul/sample) of 5, 10, 15, 20, 30, 40 and 60. No optimization is required if using same volumes. Pools were then processed and extracted as previously described. Each reaction contained the undiluted sample used for pooling to assess for sample degradation of variation between assays. Finally, 5ul of the extracted RNA was added to the RT-qPCR reagent mix from Kogene or Dyatheva. We repeated the experiment two times and all the pooled samples were ran in triplicate (Figures 1 and 2). As shown in Figure 1, all the pools were positive for RdRp gene, which is consistent with other reports selecting molecular targets for COVID-19 diagnostics (*13*). As the sample pool increased signal for E gene was lost after pooling 20 negative samples. However, Diatheva kit managed to detect both targets in the pool of 30 samples (Figure 2).

**Figure 1.**
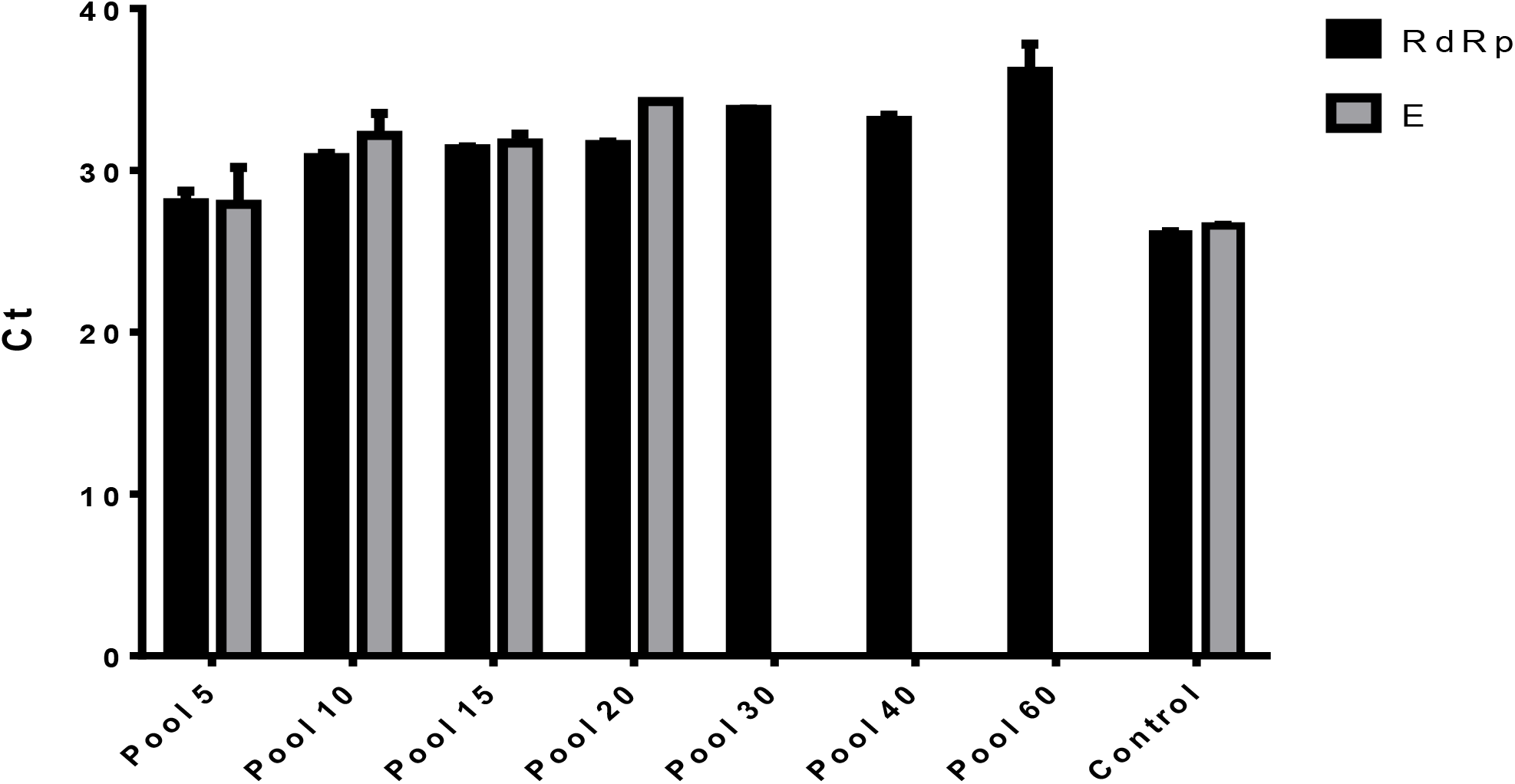
PCR results using the pools of 5, 10, 15, 20, 30, 40 and 60 using Kogene for RdRp and E genes. Results are presented as average Ct from two independent experiments.

**Figure 2.**
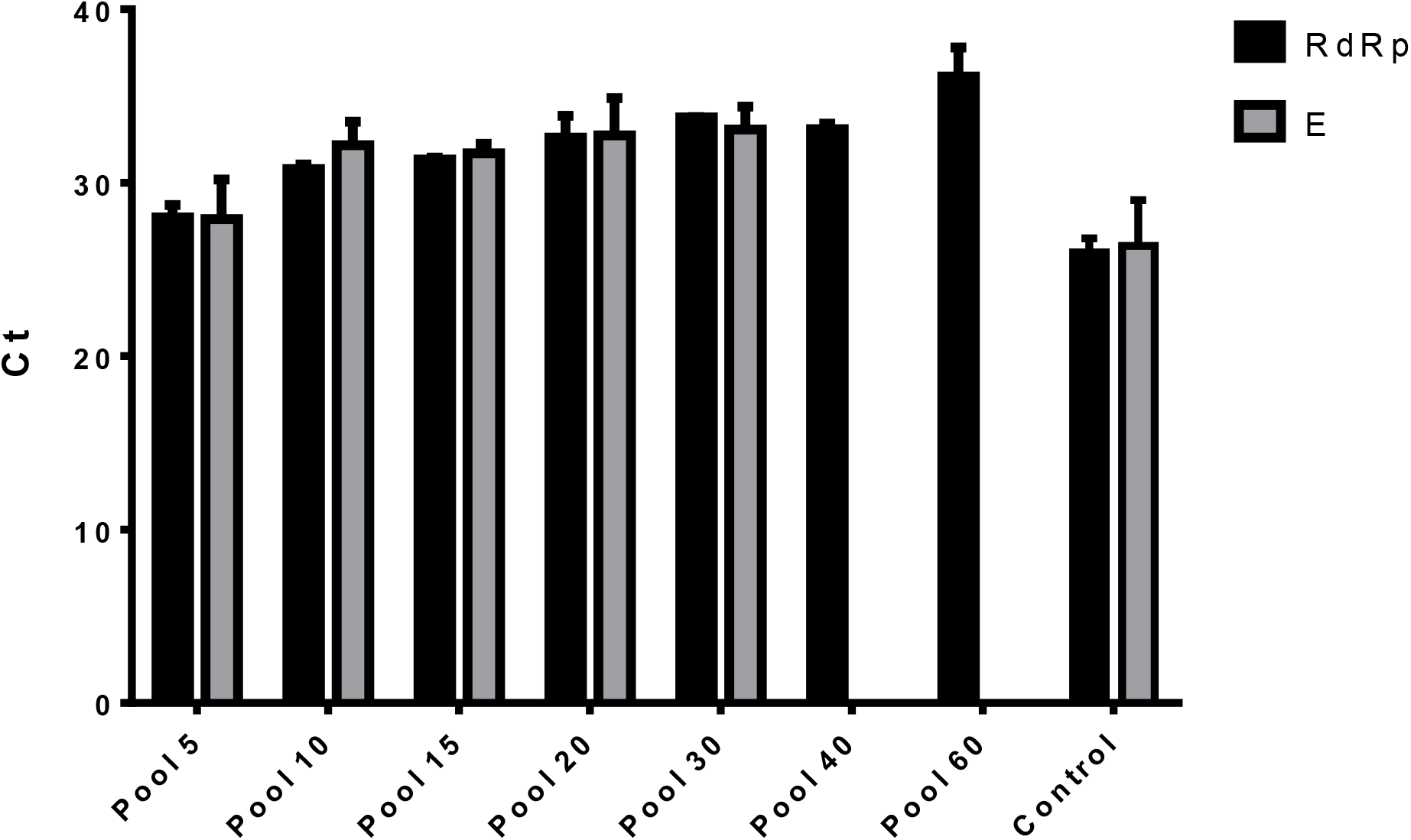
PCR results using the pools of 5, 10, 15, 20, 30, 40 and 60 using Diatheva kit for RdRp and E genes. Results are presented as average Ct from two independent experiments

## Discussion

As some countries are lifting up lockdown measures implemented through February and March, economies have to open quickly and safely. The initial steps to resume economical activities have to prioritize public health (*14*). Therefore massive scaling up COVID-19 testing is the temporary solution until immunity levels are achieved. One of the approaches that can be easily applied in order to increase the number of tests represents sample pooling. We showed that using a range of negative sample matrixes with one representative sample, ranging from 5 to 60 pools only leads to only an incremental increase in the Ct values, for the RdRp target. This is consistent with the initial report for SARS-CoV-2 pooling, however the number of samples used for pooling before RNA extraction, was limited to 7 (*12*). This approach would be feasible for laboratories that are performing large-volume testing and considering screening with the commercial kits evaluated in this study. Moreover, laboratories may consider testing as many as 60 samples using different sample matrixes, using the standard protocols, as an option for cost savings without compromising the capacity to detect SARS-CoV-2. However, there are several limitations that might arise using the pooled sample approach.

One limitation of this approach is that it seems only the RdRp gene is suitable for detection of SARS-CoV-2 in pools larger than 30 samples, which might arise in false negative results due to equipment variation and sample handling. However, this could be easily circumvented by integrating additional SARS-CoV-2 specific targets. Moreover, the complexity of the disease can influence the sensitivity and specificity of the assay (*15*). Our results are showing that RdRp would be the ideal target for sample pooling, rather than E gene. This agrees with the initial development of molecular tests for SARS-CoV-2 detection, when RdRp gene assays 3.6 copies per reaction for the RdRp assay (*4; 13*).

In this research, we showed that sample pooling for SARS-CoV-2 diagnostic is a feasible measure using commercial kits widely available.

## Competing Interest Statement

The authors have declared no competing interest.

## References

1. World Health Organization (WHO). Coronavirus. Geneva: WHO; 2020 [Accessed 25 May 2020]. Available from: https://www.who.int/health-topics/coronavirus

2. Zhu, N. et al. A novel coronavirus from patients with pneumonia in China, 2019. N. Engl. J. Med. 382, 727–733 (2020).

3. Esbin MN, Whitney ON, Chong S, Maurer A, Darzacq X, Tjian R. Overcoming the bottleneck to widespread testing: A rapid review of nucleic acid testing approaches for COVID-19 detection [published online ahead of print, 2020 May 1]. RNA. 2020; rna.076232.120. doi:10.1261/rna.076232.120

4. Corman VM, Landt O, Kaiser M, Molenkamp R, Meijer A, Chu DK, Bleicker T, Brünink S, Schneider J, Schmidt ML, Mulders DG, Haagmans BL, van der Veer B, van den Brink S, Wijsman L, Goderski G, Romette JL, Ellis J, Zambon M, Peiris M, Goossens H, Reusken C, Koopmans MP, Drosten C. 2020. Detection of 2019 novel coronavirus (2019-nCoV) by real-time RT-PCR. Euro Surveill 25(3). doi:10.2807/1560-7917.ES.2020.25.3.2000045.

5. Chu DKW, Pan Y, Cheng SMS, et al. Molecular Diagnosis of a Novel Coronavirus (2019-nCoV) Causing an Outbreak of Pneumonia. Clinical Chemistry. 2020 Apr;66(4):549–555. DOI: 10.1093/clinchem/hvaa029.

6. Wölfel, R., Corman, V.M., Guggemos, W. et al. Virological assessment of hospitalized patients with COVID-2019. Nature 581, 465–469 (2020). https://doi.org/10.1038/s41586-020-2196

7. Detmer, S.E., Patnayak, D.P., Jiang, Y., Gramer, M.R., Goyal, S.M., 2011. Detection of Influenza a Virus in Porcine Oral Fluid Samples. J VET Diagn Invest 23, 241–247. doi:10.1177/104063871102300207

8. Prickett, J.R., Zimmerman, J.J., 2010. The development of oral fluid-based diagnostics and applications in veterinary medicine. Anim Health Res Rev 11, 207–216. doi:10.1017/S1466252310000010

9. Grau, F.R., Schroeder, M.E., Mulhern, E.L., McIntosh, M.T., Bounpheng, M.A., 2015. Detection of African swine fever, classical swine fever, and foot-and-mouth disease viruses in swine oral fluids by multiplex reverse transcription real-time polymerase chain reaction. J. Vet. Diagn. Invest. 27, 140–149. doi:10.1177/1040638715574768

10. Sullivan TJ, Patel P, Hutchinson A, Ethridge SF, Parker MM. Evaluation of pooling strategies for acute HIV-1 infection screening using nucleic acid amplification testing. J Clin Microbiol. 2011;49(10):3667‐3668. doi:10.1128/JCM.00650-11

11. Nguyen, N.T., Aprahamian, H., Bish, E.K. et al. A methodology for deriving the sensitivity of pooled testing, based on viral load progression and pooling dilution. J Transl Med 17, 252 (2019). https://doi.org/10.1186/s12967-019-1992-2

12. Yelin I, Aharony N, Shaer Tamar E, et al. Evaluation of COVID-19 RT-qPCR test in multi-sample pools [published online ahead of print, 2020 May 2]. Clin Infect Dis. 2020; ciaa531. doi:10.1093/cid/ciaa531

13. Jasper Fuk-Woo Chan, Cyril Chik-Yan Yip, Kelvin Kai-Wang To, Tommy Hing-Cheung Tang, Sally Cheuk-Ying Wong, Kit-Hang Leung, Agnes Yim-Fong Fung, Anthony Chin-Ki Ng, Zijiao Zou, Hoi-Wah Tsoi, Garnet Kwan-Yue Choi, Anthony Raymond Tam, Vincent Chi-Chung Cheng, Kwok-Hung Chan, Owen Tak-Yin Tsang, Kwok-Yung Yuen. Journal of Clinical Microbiology Apr 2020, 58 (5) e00310–20; DOI: 10.1128/JCM.00310-20

14. https://www.forbes.com/sites/kenrapoza/2020/04/20/european-economies-to-open-before-the-us/

15. Long, C., Xu, H., Shen, Q., Zhang, X., Fan, B., Wang, C., Zeng, B., Li, Z., Li, X., & Li, H. (2020). Diagnosis of the Coronavirus disease (COVID-19): rRT-PCR or CT?. European journal of radiology, 126, 108961. https://doi.org/10.1016/j.ejrad.2020.108961

